# Song as a behavioural pre-mating barrier in early population divergence: Insights from the Canary Islands Chaffinch

**DOI:** 10.64898/2026.03.12.711316

**Authors:** Bárbara Freitas, Diego Gil, Christophe Thébaud, Borja Milá

## Abstract

Acoustic signaling is key to individual and species recognition, playing a major role in sexual and social communication. Since reproductive isolation is often maintained through pre-mating mechanisms, song can be an early isolating trait leading to assortative mating, promoting reproductive divergence, and potentially contributing to speciation. However, whether song differences alone are sufficient to prevent interbreeding or if other traits also contribute, remains a matter of debate. Playback experiments provide a more direct way to test the role of song as a reproductive barrier. Here, we use playback experiments to test the hypothesis that song acts as a pre-mating barrier in two recently diverged populations of an island passerine, the Canary Islands Chaffinch (*Fringilla canariensis palmae*), which inhabit ecologically distinct laurel and pine forests within the island of La Palma. Assuming that male song has diverged in the two habitats, we tested if territorial males from a given habitat responded differently to songs from intruding males from their own habitat or from the other habitat type, using a closely related mainland species as a control. We found that probability of response was weaker to songs of the closely related species and to the different-habitat birds than to songs of the same-habitat birds, but differences for the latter were weak. The intensity of response followed the same pattern. Overall, song divergence between laurel and pine forest chaffinches does not appear strong enough to cause clear behavioural discrimination against individuals from the alternative habitat. Other factors such as morphological and ecological divergence associated with adaptation to local resources might better explain population differentiation. However, testing female responses will be essential to determine whether songs convey lineage-specific information that may elicit assortative mating.

---

In birds, reproductive isolation is thought to be primarily maintained by pre-mating isolation mechanisms and only secondarily by post-mating incompatibilities (Grant & Grant, 1997; Edwards *et al*., 2005). Acoustic signalling, which plays a major role in individual and species recognition (Gil & Gahr, 2002; Catchpole & Slater, 2008), is often hypothesised to be implicated in the first stages of the development of reproductive isolation. Differences in song can prevent populations from recognising each other as conspecifics, which may lead to assortative mating and contribute to the build-up of reproductive isolation, and potentially, speciation (Irwin & Price, 1999; Slabbekoorn & Smith, 2002; Uy, Moyle & Filardi, 2009; Freeman & Montgomery, 2017).

Consequently, quantifying acoustic divergence between populations may appear to provide a straightforward way to assess whether recognition systems have diverged sufficiently to reduce mating opportunities. However, a potential limitation of assessing vocal divergence through the analysis of acoustic traits is that the vocal cues that can be measured might not be those actually used by birds in mating decisions. Thus, even if populations show differences in acoustic traits, these differences may not be biologically relevant or sufficient to prevent interbreeding (Nelson, 1998; Soha *et al*., 2016).

Experimental approaches can help understand the relationship between acoustic traits and reproductive isolation, with playback experiments providing an effective way to directly assess how birds respond to the songs of individuals from their own population relative to those of individuals from other populations. These experiments attempt to “ask the birds themselves” whether they perceive vocal differences as biologically relevant, providing a useful proxy for premating reproductive isolation (Soha *et al*., 2016; Freeman & Montgomery, 2017). Playback responses are typically measured through territorial behaviour, because territory defence in both inter- and intraspecific contexts is often mediated by vocal signals, allowing individuals to identify and respond to competitors (e.g., Gil, 1997). Effective territory defence depends on recognising signals at a distance and distinguishing the species, sex, and identity of the signaller, while also assessing relevant attributes such as individual quality and motivation (Catchpole & Slater, 2008; Niśkiewicz, *et al*., 2024). Thus, for song differences to play a role in the early evolution of reproductive isolation, individuals must either fail to recognise or at least respond less aggressively to foreign acoustic signals than to their own.

Studies on song discrimination have produced mixed results on whether song divergence may initiate assortative mating and lead to reproductive isolation. In a number of cases, birds responded more strongly to playbacks of songs from their own population than to songs from a different population (Salomon, 1989; Liu *et al*., 2008; Reichard, 2014; Freeman & Montgomery, 2017), which suggests that song may play a role in discrimination between lineages that have diverged but are not yet reproductively isolated, and thus could contribute to the early build-up of reproductive isolation if such discrimination reduces mating opportunities between these lineages. However, this is not always the case, and in other studies, birds responded similarly to the songs of both local and foreign populations (Irwin, Bensch & Price, 2001; Dingle *et al*., 2010), suggesting that song divergence alone may not always contribute to reproductive isolation, at least in its early stages.

In species in which discrimination occurs, males usually sing relatively stereotyped simple songs, where discrete changes in song sub-units (i.e., syllables) can delimit song boundaries across populations (see Gallego-Abenza *et al*., 2024 for examples). In turn, it has been argued that greater song complexity or larger repertoires may limit the ability to discriminate between similar songs, eliciting non-specific responses in receivers (Kroodsma, 1976; Catchpole & Slater, 2008), yet recent research using laboratory populations of zebra finches has shown that discrimination of subtle acoustic variation in vocal signals is possible in songbirds with complex learnt songs (Fishbein *et al*., 2021).

Here, we assessed the importance of song as a pre-mating barrier in early population divergence by focusing on the Canary Islands Chaffinch, a species with a complex learnt song, on the island of La Palma (*Fringilla canariensis palmae*). Ancestors of this species colonised the laurel forests of La Palma within the last 100 000 years, and subsequently spread to the extensive dry pine forests that cover most of the island (Recuerda *et al*., 2021, 2023). La Palma’s contrasting habitats - humid laurel forests (hereafter “cloud forest”) and dry pine forests (hereafter “pine forest”) - present distinct ecological conditions, characterised by differences in plant species composition, vegetation structure, food resources and climatic conditions (Irl & Beierkuhnlein, 2011). Recent findings by Recuerda *et al*. (2023) show evidence of adaptive phenotypic and genomic divergence between populations from these forest types despite the small geographic scale involved. Birds in pine and cloud forests differed in stable isotope signatures (δ¹³C, δ¹⁵N), morphology, and plumage coloration in ways that are consistent with ecological and geographical predictions. Moreover, genome-wide analyses revealed both neutral genetic structure, reflecting low natal dispersal, and adaptive divergence at loci associated with phenotypic and environmental variation (Recuerda *et al*., 2023). Together, these results suggest the role of ongoing local adaptation and potentially sexual selection in driving population differentiation. Given this ecological divergence and genetic structure, this system provides an ideal context to test whether song discrimination may be acting as a behavioural isolation mechanism between recently diverged populations of *F. c. palmae*.

In this study, we test for (i) the probability of an aggressive response (i.e., whether a response to the stimulus occurs), and (ii) the intensity of the aggressive response (i.e., the strength of the aggressive reaction). We hypothesise that vocal differences between populations in these adjacent habitats may restrict mating opportunities between them, driving population divergence if responses to the same-habitat songs are stronger and more frequent relative to different-habitat songs and to songs from the closely related Common Chaffinch (*Fringilla coelebs*). Such responses would indicate that chaffinch populations from the two habitats of La Palma recognise members of their own habitat as more direct sexual and/or ecological competitors and that song may be involved in male-male competition and potentially female choice (Baker & Baker, 1990; Baker, 1991; Patten, Rotenberry & Zuk, 2004). In contrast, a similarly aggressive response to same- and different-habitat stimuli would suggest that male chaffinches do not discriminate between songs from cloud and pine forest habitats, instead perceiving males from both forest types as equally concerning competitors. If we assume that male aggressive behaviour is proportional to female sexual attraction towards males from the same and different habitat, then this result would mean that song does not represent a major reproductive barrier at this early stage of divergence.

## METHODS

### Study areas and song sampling

The study was conducted in sites located in the two forest types on La Palma island: six localities of cloud forest and five of pine forest (Fig. 1A). The vegetation communities of these forests are shaped by the trade winds, which on La Palma blow in a NE to SW direction, so that cloud forests are present in mid-elevations along the northern and eastern slopes and are dominated by evergreen trees from the Lauraceae family (Fernández-Palacios, 2018). In contrast, pine forests extend between 500 m a.s.l. on the leeward side and 1100-2000 m a.s.l. on the windward side and consist of open dry woodland dominated by the endemic Canary pine (*Pinus canariensis*). Mean annual precipitation ranges from 500 to 1200 mm in the cloud forest and from 450 to 600 mm in the pine forest. Mean annual temperature ranges between 13 and 18°C in cloud forest and between 11 to 15°C in pine forest (del Arco Aguilar & Rodríguez Delgado, 2018).

**Fig. 1.**
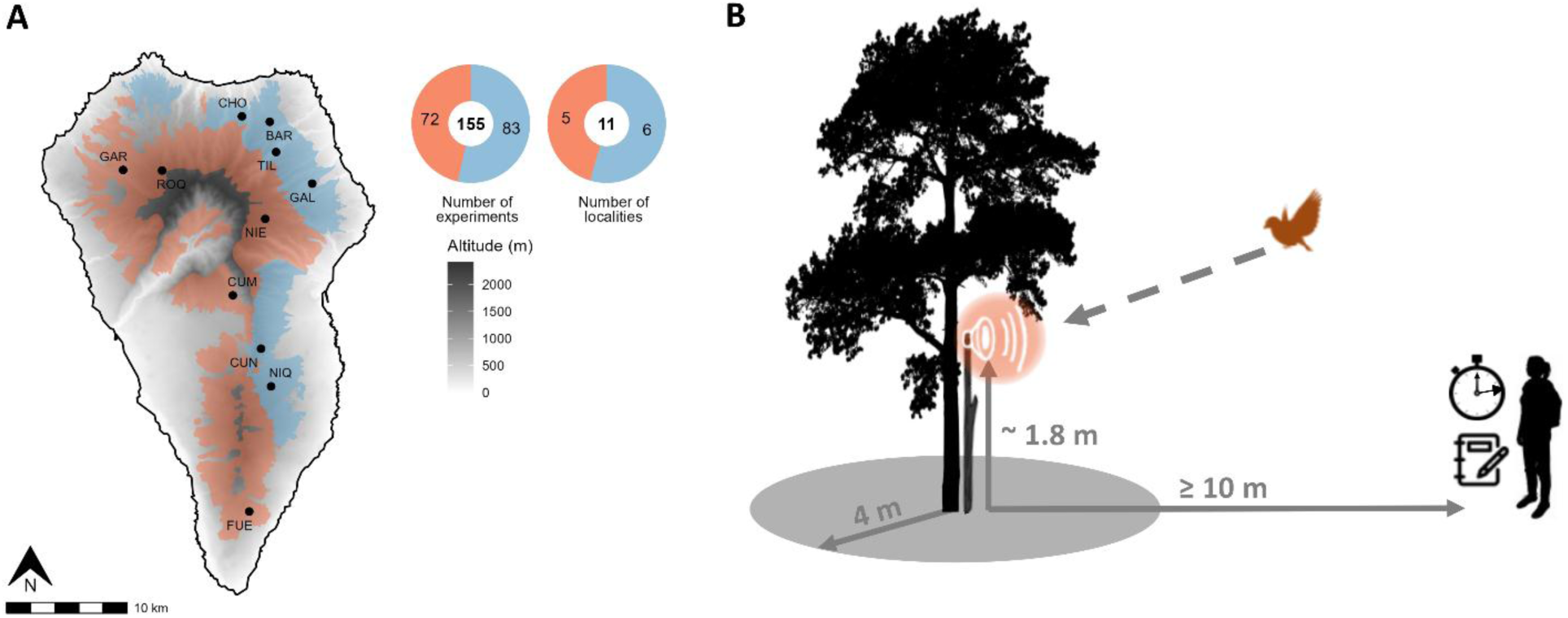
A) Map of La Palma island, showing the distribution of pine forest (orange) and cloud forest (blue) and the localities where the playback experiments were conducted. Doughnut charts show the number of experiments and the number of localities sampled in each forest type (totals in the centre); BAR = Barlovento, CHO = Charca de Ovejas, CUM = Cumbrecita, CUN = Cumbre Nueva, FUE = Fuencaliente, GAL = La Galga, GAR = Garafía, NIE = Pico de las Nieves, NIQ = Niquiomo, ROQ = Roque de los Muchachos, TIL = Los Tilos, B) Playback experiment setting.

Recordings of male birds were taken in May 2021, April-May 2022, and April-May 2023 during the morning hours (0650 - 1155 h, earliest sunrise at 0717 h verified at http://timeanddate.com), aiming for approximately two-minute recordings per individual using a Marantz PMD-670 audio recorder (Marantz Professional, Kanagawa, Japan) equipped with a Sennheiser ME67 shotgun microphone (Sennheiser, Wedemark, Germany), or a SoundDevices 722 audio recorder with a Sennheiser MKH 70 P48 microphone, or a Tascam DR-40X digital recorder with an external Sennheiser ME66 microphone. Recordings were saved as 16-bit WAV files at a sampling frequency of 44 and 48 kHz.

Recordings of the Common Chaffinch were taken during the breeding season of 2021 (specifically March, April, and June), at Soto del Real, Spain, during morning hours (0700 - 1300 h, earliest sunrise at 0643 h).

### Playback stimulus songs

We prepared four types of playback stimuli using Audacity (Audacity Team, 2021): (i) chaffinch songs from La Palma’s pine forest, (ii) chaffinch songs from La Palma’s cloud forest, (ii) Common Chaffinch songs from mainland Spain, and (iv) white noise to be used as a control. For the first three stimulus types, we selected songs with a good signal-to-noise ratio from individuals previously recorded across different localities. For the playback experiments performed in 2022 (see ‘Playback experiment design and behavioural responses’), we prepared three stimuli per type of playback stimulus using songs from different individuals for each playback, except for the Common Chaffinch playback, for which we prepared only two stimuli. As we expanded our dataset of individuals recorded from each locality in La Palma, we incorporated additional recordings to the playback experiments performed in 2023 to include songs from a total of nine individuals from the cloud habitat and ten from the pine habitat (Table S1).

Each playback stimulus consisted of a single song followed by a 4-second silent period (approximating the natural calling rate of the Common Chaffinch in La Palma), repeated over a 1-minute period. Each stimulus song was selected from a different individual, aiming to avoid pseudo-replication while maintaining a broadly comparable song structure. Each recording was filtered with a high-pass filter of 0.9 kHz and low-pass filter of 9 kHz, normalised (−24 db), and saved as an uncompressed 16-bit wave file (.wav). For the white noise stimulus, we created periods of white noise with the amplitude set at 0.8 (default) and alternated them with 4-second silence periods for a total period of 1 minute. We created three tracks of white noise stimulus, in which we defined the duration of the white noise periods as 2.4, 2.6, and 2.8 seconds, respectively. As the intensity of territorial behaviour in chaffinches is modulated by the amplitude of songs to which they are exposed (Brumm & Ritschard, 2011), we calibrated the amplitude of each playback and set it at about 82 dB SPL measured at a 1-m distance from the speaker, using a CESVA SC-2 digital sound level meter (CESVA instruments s.l.u, Barcelona, Spain). This amplitude is the mean value of free-ranging male chaffinches measured in their natural habitat by Brumm & Ritschard (2011). Detailed information on the original recordings used for playback stimuli is available in Table S1.

### Playback experiment design and behavioural responses

Playback experiments took place during the breeding season (7^th^ April – 10^th^ May 2022 and 20^th^ April – 21^st^ May 2023), between 0600 and 1130 h (earliest sunrise at 0717 h verified at https://www.timeanddate.com/). The time of testing reflected the birds’ period of maximum activity (Cramp, Perrins & Brooks, 1994) and depended on the weather on any particular day. Experiments were conducted only in areas where territorial individuals had been previously confirmed (either captured or heard/recorded). Playback points were set at least 150 m apart from each other to prevent the possibility of testing the same individual more than once, a distance considered sufficient given personal observation and consistent with reported population densities for the Common Chaffinch of 13–161 individuals/km² in April-May (Santini, Isaac & Ficetola, 2018). Before each trial, we placed a small speaker (JBL Clip 4, dynamic frequency response of 100 Hz – 20 kHz; Harman International Industries, Stamford, Connecticut, USA) at a height of approximately 1.8 m, near a bush or branch that an approaching bird could use as a perch (Fig. 1B). In cases where this was not possible due to absence of middle layer forest (most frequently in pine forest), we attached the speaker to a pole and placed it close to the nearest tree. We never tested neighbours with the same playback stimulus. The playback stimuli were played in a counterbalanced order and from a mobile phone via Bluetooth connection to the speaker. Although Bluetooth transmission involves lossy compression, potentially reducing signal fidelity in comparison to wired connections, the discrimination responses detected across treatments, particularly for the Common Chaffinch stimuli (see Results), suggest that any reduction in signal fidelity did not prevent effective discrimination, and it may have even resulted in more conservative estimates of behavioural differences. While we were unaware of the potential implications of Bluetooth compression at the time of the study, we recognise the importance of this issue for future playback designs and recommend caution and further assessment of its impact on non-human auditory perception.

Playback sessions consisted in one of the following treatments: (i) same-habitat (songs from La Palma’s pine forest played to pine forest individuals, and songs from the cloud forest played to cloud forest individuals); (ii) different-habitat (pine-to-cloud and cloud-to-pine); (iii) Common Chaffinch; and (iv) control (white noise). Each playback session lasted 10 minutes: 1 minute of broadcast stimulus, followed by a 9-minute observation period without playback. Behavioural data collection started when the first song of the stimulus was broadcast and was taken from about 10 m distance from the speaker, to avoid missing bird movements, especially in cloud forest. To document individual responses, we took notes manually and continuously. If a response was elicited from more than one individual in the area, we monitored only the first individual with whom we established eye contact. We assessed playback responses by measuring parameters based on previous playback experiments with species of the same genus (Slater & Catchpole, 1990; Leitão & Riebel, 2003): (1) time to approach the speaker; (2) closest approach to the speaker; and (3) time spent within 4 m of the speaker. Approach to the speaker has been widely interpreted as an aggressive signal and is considered a valid measure of territorial response strength in chaffinches (Marler, 1956; Hinde, 1958; Slater & Catchpole, 1990; Vallet & Kreutzer, 1992; Naguib *et al*., 2000) and in other songbird genera (e.g., Vehrencamp, 2001; Molles & Vehrencamp, 2001). Singing by the territorial male in response to the stimulus was also noted, but it was too rare for statistical analysis. Chaffinches do not generally respond to playback with song (Slater, 1981). When no evidence of approach was obtained, the latency to approach was set at the maximum time interval of 10 minutes and the closest approach was scored as 100 m. These experiments were categorised as ‘no response’ compared to ‘response’ experiments, in which at least one of the three playback parameters was measured. Wind speed was also registered according to an adapted Beaufort scale (0 = no wind, 1 = grass is moving, 2 = small branches of trees are moving, 3 = large branches of trees are moving, 4 = the trunks are moving; European Stag Beetle Monitoring Network, 2024).

All birds were only tested once and resumed normal activity (were observed foraging) after the experiment. Distance measures of responding birds were assessed by comparison with the height of the speaker, whose distance from the ground was known, and time measures were recorded with a stopwatch running for 10 min during each test. To ensure consistency, all behavioural observations and transcription of data were conducted by the same person (BF).

### Ethical note

To our knowledge, the individual birds tested in the experiment reflected the population in a representative way with no apparent biases resulting from social background, self-selection, habituation, or other factors as recommended in the STRANGE framework (Webster & Rutz, 2020). Our experimental procedure adhered to the ASAB/ABS Guidelines for the care and use of animals (Asab Ethical Committee/ABS Animal Care Committee, 2023). Field work in La Palma was conducted under research permits A/EST-004/2020 and A/EST-008/2023.

### Statistical Analysis

In our experimental design, the white-noise treatment was intended as a negative control, so that we expected birds to not respond aggressively to the speaker when a white-noise was emitted. This was confirmed (i.e., no responses were observed for the control playback treatment; Fig. 2, Table S2), so this treatment was not included in further analyses.

**Fig. 2.**
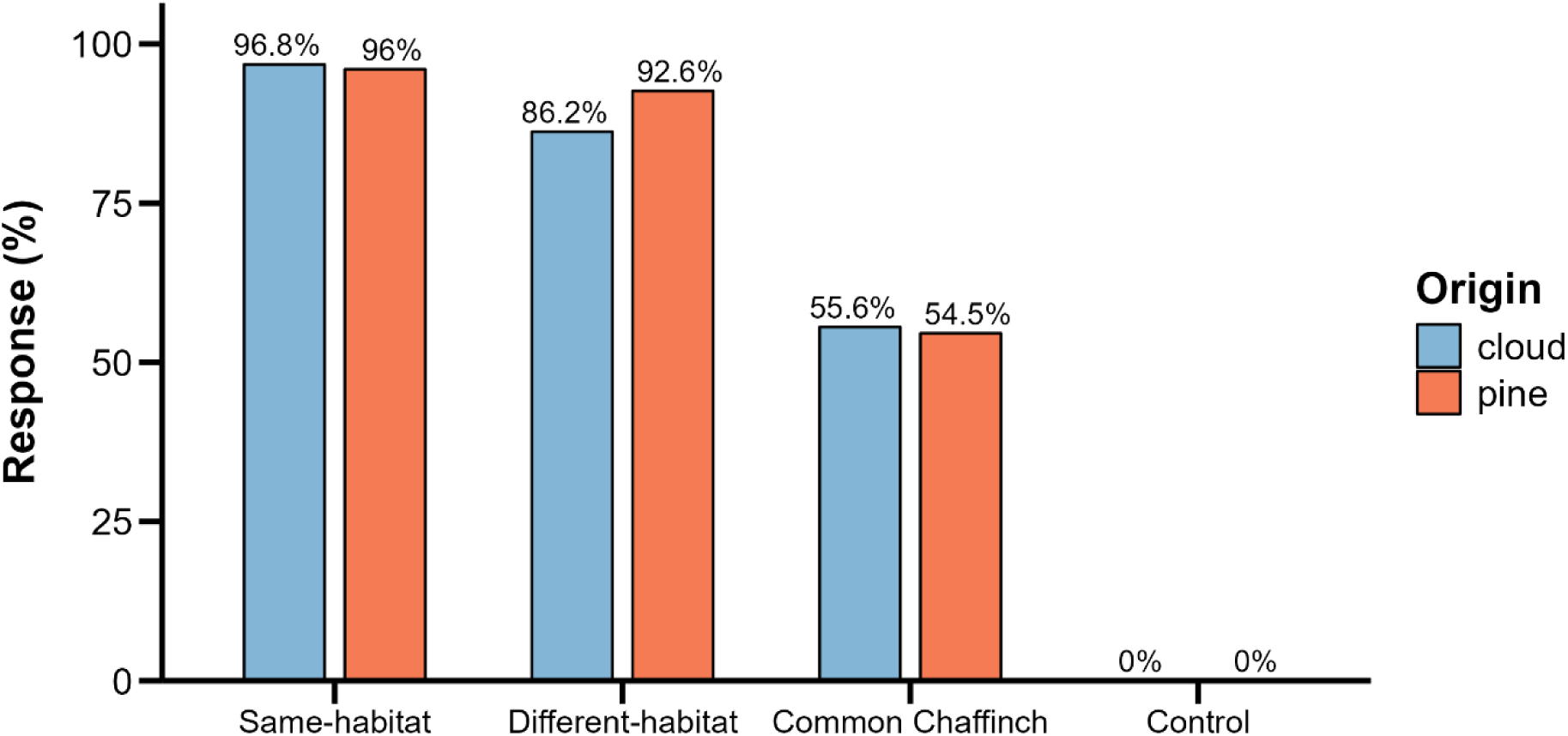
Percentage of responses to the playback experiments for each treatment category (same-habitat, different-habitat, Common Chaffinch, and control), presented separately for each habitat type (origin). Total number of experiments performed by order (left to right): 31, 25, 29, 27, 9, 11, 14, 9.

**Fig. 3.**
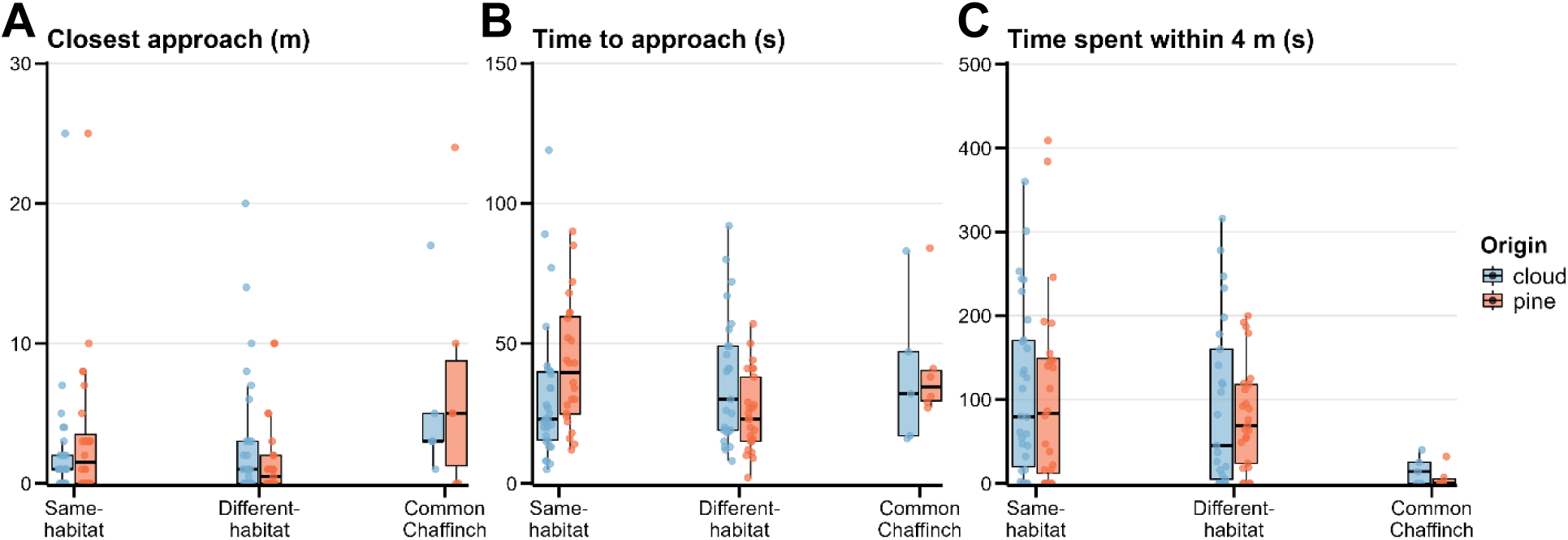
Plots of the behavioural variables registered during the playback experiments, depending on the experimental treatment (same-habitat, different-habitat, and Common Chaffinch), showing the median, upper and lower quartiles, whiskers, and points representing the response for each individual adult male.

Our aim was to use the behavioural response to playback experiments as a proxy for behavioural isolation based on song. Thus, we define ‘‘song discrimination’’ here as a territory owner responding to same-habitat song more often and more aggressively than to the other treatments. For this, we distinguished between responsiveness probability (‘‘response’’ versus ‘‘no response”) and intensity of the response (‘‘weak’’ versus ‘‘strong’’ responses).

### Responsiveness probability

To assess the probability of an aggressive response during playback experiments, we fitted a global generalised linear mixed-effects model (GLMM), with a binomial error distribution and logit link function, using the ‘glmer’ function from the ‘lme4’ R package (Bates *et al*., 2015). The response variable was a binary measure indicating whether an individual responded (‘1’) or did not respond (‘0’) to the playback. The fixed effects in the model included the categorical playback treatment and the type of habitat where we performed the experiment (‘origin’). Origin was included in the model because the distinct vegetation structures of the forest types could influence perch availability or suitable locations for birds to approach, potentially impacting the recorded behavioural responses. We also included as fixed effects the year, wind scale, minutes after sunrise and the day of the year (ISO_8601 scale), as birds may respond differently to territorial intrusions throughout the day and season. We included locality ID as random effects to account for variability in responses across individuals of different localities. Minutes after sunrise and the day of the year were centred and scaled prior to analysis to improve model convergence. Models were checked for convergence and singular fits before model selection. After fitting the full model, we computed all possible model combinations (but always keeping the random factor and playback treatment and origin as fixed factors) and ranked them based on the Akaike Information Criterion for small sample sizes (AICc). All models within ΔAICc ≤ 2 of the top-ranked model were retained and parameter estimates were obtained using model averaging. Model assumptions were evaluated using simulation-based residual diagnostics of the global model using the ‘DHARMa’ package (Hartig, 2024).

### Intensity of the response

To assess whether individuals responded more strongly to playbacks from the same habitat, a different habitat, or to the Common Chaffinch, we analysed each behavioural response variable (time spent within 4 m of the speaker, closest approach distance, and time to approach the speaker) separately. These analyses were restricted to individuals that responded to the playback. For time spent within 4 m of the speaker, we fitted a linear mixed-effects model with the same fixed and random effects structure as described above for responsiveness probability: playback treatment, origin, year, wind scale, minutes after sunrise, and day of the year (ISO 8601 scale) were included as fixed effects, and locality ID as a random effect. Model assumptions were evaluated using simulation-based residual diagnostics with the DHARMa package (Hartig, 2024; Fig. S1). For time to approach and closest approach, the random effect of locality ID explained negligible variance and resulted in singular fits. Therefore, locality ID was removed and the response was analysed using linear fixed-effects models only.

For each response variable, we first fitted a global model including all fixed effects. We then conducted model selection using the ‘dredge’ function in the ‘MuMIn’ R package (Barton, 2022), ranking all candidate models based on AICc while retaining treatment and origin in all models. All models within ΔAICc ≤ 2 of the top-ranked model were retained and parameter estimates were obtained using model averaging. Closest approach and time to approach were right-skewed, so we log-transformed the response variable (log(x + 0.01)). Model assumptions were evaluated using simulation-based residual diagnostics of the global model using the ‘DHARMa’ package (Hartig, 2024; Fig. S1-3). All measures are presented as estimate ± adjusted standard error, unless otherwise indicated.

## RESULTS

In total, we performed 155 playback experiment sessions, of which 56 were same-habitat (31 in cloud and 25 in pine forest), 56 were different-habitat (29 in cloud and 27 in pine forest), 20 were Common Chaffinch (9 in cloud and 11 in pine forest), and 23 were controls (14 in cloud and 9 in pine forest; Fig. 2 and Table S2). These playback experiment sessions were performed in 11 different localities across La Palma, 6 in cloud forest and 5 in pine forest (Fig. 1A).

### Responsiveness probability

Canary Islands Chaffinch males were significantly less likely to respond to Common Chaffinch songs than to songs of conspecifics from either habitat within La Palma (β = −3.375 ± 0.968, *P* < 0.001). Responses to different-habitat playbacks were also less likely compared to same-habitat playback, but this difference was weak and statistically non-significant (β = −1.250 ± 0.886, *P* = 0.158). The probability of response did not significantly differ between individuals from pine compared to cloud forest populations (origin; β = 0.084 ± 0.830, *P* = 0.920), suggesting that habitat origin did not influence the pattern of response to different playback types (Table 1). The probability of response tended to be lower in 2023 than in 2022, although this effect was weak (β = -0.916 ± 0.883, *P* = 0.300).

**Table 1.** Summary of the generalised linear mixed-effects model (GLMM) full average results for the responsiveness probability. Fixed effects include treatment and origin. Random effects account for variance among populations (Population ID). Estimates (β) are presented with their adjusted standard error (Adjusted SE), z-values, and P-values. Significant effects (*P* < 0.05) are indicated in bold.

| Effect | Estimate ($\beta$) | Adjusted SE | z-value | P-value |
| --- | --- | --- | --- | --- |
| Intercept | 3.962 | 1.010 | 3.922 | **<0.001** |
| Treatment: different-habitat | -1.250 | 0.886 | 1.411 | 0.158 |
| Treatment: Common Chaffinch | -3.375 | 0.968 | 3.488 | **<0.001** |
| Origin: pine | 0.084 | 0.830 | 0.101 | 0.920 |
| Year: 2023 | -0.916 | 0.883 | 1.038 | 0.300 |
| Effect | Variance ($\sigma^2$) | SD | | |
| Population ID | 0.360 | 0.599 |  |  |

### Intensity of the response

Males spent significantly less time within 4 m of the speaker during playbacks of the Common Chaffinch compared to same-habitat songs (β = −98.532 ± 29.947, *P* = 0.001; Table S3). Males also spent less time within 4 m during playbacks of the different-habitat compared to same-habitat songs (β = -19.160 ± 17.580, *P* = 0.276), but this trend was weak. No significant effects were detected for origin, day of the year, year, or wind scale (Table S3).

Although there was a pattern of gradually less intense responses across treatments (same-habitat, different-habitat, or Common Chaffinch; Table S4), closest-approach distance did not differ significantly among treatments. Birds approached significantly less closely in 2023 than in 2022 (β = - 1.423 ± 0.562, *P* = 0.011).

Time to approach the speaker did not differ significantly among treatments (same-habitat, different-habitat, or Common Chaffinch; Table S5), indicating no differences on how quickly males approached the speaker. Similarly, origin, wind scale, and minutes after sunrise had no significant effect on time to approach.

## DISCUSSION

Song is often hypothesised to be an early isolating mechanism that can promote assortative mating, and potentially contribute to population divergence and speciation. However, there is ongoing debate about whether song differences alone are sufficient to prevent interbreeding during early stages of speciation. We performed playback experiment sessions across different localities in La Palma to determine if song could act as a biologically relevant cue for isolation between populations of Canary Islands Chaffinches that have recently diverged in two different habitat types on La Palma. Our results showed that male chaffinches from both cloud and pine forests in La Palma responded less often and less strongly to song playbacks of the closely related Common Chaffinch from the mainland, indicating species-level discrimination. Territorial males also replied more frequently and more strongly to songs from conspecifics of the same-habitat than of different-habitat, but this effect was weak and not statistically significant, suggesting that, while the differences in this trait might not yet be strong enough to promote behavioural isolation, they may represent an intermediate step in that direction. We discuss these findings below, while also identifying limitations of our study and proposing directions for future research.

### Limited song discrimination

The playback experiment showed that male chaffinches from both habitat types in La Palma were more likely to reply and to reply more strongly to males from the same habitat than to males from the different habitat, although in our analyses this differential aggressive response was weak and not statistically significant. If song was a key isolating mechanism, we would expect birds to show a stronger response to songs from their own habitat, but this pattern was not observed. Thus, our results provide weak, non-conclusive support for our hypothesis that song divergence between populations in different habitats contributes to behavioural isolation in this system. Island birds have been reported to have songs with greater syllable diversity relative to continental counterparts, possibly as a result of reduced interspecific competition on islands (Kroodsma, 1985; Robert *et al*., 2022). This greater diversity could make it more difficult for individuals to discriminate between songs from different populations, and would explain the lack or low levels of song discrimination in the Canary Island Chaffinch from La Palma.

Previous studies on closely related species are relatively consistent with our results. For instance, the experiment by Slater & Catchpole (1990) in Tenerife island, found that the local subspecies *F. c. canariensis* responded to songs of the mainland species, *Fringilla coelebs gengleri*, nearly as often as to their own, although the response was less intense. Similarly, no significant differences were found in the response of the Blue Chaffinch *Fringilla teydea* to both *F. c. canariensis* and *F. c. gengleri* songs.

However, some limitations of the playback experiments should be considered when interpreting these results. Although we selected variables known to be reliable indicators of behavioural response to playback in previous studies, variation in latency to respond may have been influenced by the bird’s initial distance from the speaker when playback began. This could introduce important levels of noise into the data, complicating the estimation of response intensity. Additionally, the observed variation in responses between years suggests that motivation or unmeasured environmental factors may influence behavioural reactions, highlighting the importance of conducting playback experiments across multiple years to assess the consistency and robustness of the observed patterns.

Additionally, because different individuals were tested for each playback treatment instead of exposing the same individuals to all treatments, variation in response may partly reflect individual differences, such as personality traits, rather than habitat-specific song recognition. Although we conducted a relatively large number of playback experiments to reduce this potential bias, individual-level variability cannot be entirely excluded.

Future improvements to playback experiments could include incorporating a visual stimulus, such as a three-dimensional model of a chaffinch with habitat-specific characteristics. This approach might better simulate a realistic territorial intrusion and elicit more accurate behavioural responses by more closely reflecting real-life scenarios.

### What male response can tell about female choosiness

Our playback experiments assessed territorial defence exerted by individual males, which is likely to be a conservative proxy for inferring the strength of a behavioural barrier to reproduction (Freeman & Montgomery, 2017). This is because for song to act as a reproductive barrier it must influence mate choice by females, and we expect females to be more selective and discerning than males, as mistakes in song recognition could lead to hybridisation and potentially maladaptive hybrid phenotypes (Qvarnström *et al*., 2006; Uy, Irwin & Webster, 2018). As a result, the response to song is expected to be broad for male birds engaging in territorial defence and mate guarding, and more narrow for females choosing mates (Seddon & Tobias, 2010). In fact, a literature review on the role of sexual learning in speciation showed that males were never found to be more selective than females (Irwin & Price, 1999).

Consequently, if a territorial bird ignores a song in a territorial context, it is likely that a female would also discriminate against that song in a mate choice context. However, if a territorial male reacts aggressively towards an intruding male of a different species, this does not mean that females would readily mate with that species, as it is adaptive for females to be more selective in choosing mates than for males to eject intruders.

A key next step would be to test female responses to playback, particularly whether females show stronger or more selective responses to songs from males of the same habitat than to those from a different habitat. Comparing the responses of both sexes would provide a more complete understanding of the role of song in reproductive isolation. For instance, Leitão and Riebel (2003) showed that although females of *F. coelebs* had preferred songs with a relatively longer flourish, males showed the strongest reaction (in that case, closest approach to the speaker) to songs with longer trills and shorter flourishes, suggesting that males and females may differently perceive aspects of chaffinch song structure and respond accordingly. However, only a limited number of studies have evaluated responses from both males and females, as collecting data on female responses can be particularly challenging. In our experiments, females rarely approached the speaker (pers. obs.), suggesting that gathering an adequate sample of female responses remains difficult in field conditions.

### Implications for speciation

The stronger response by *Fringilla canariensis palmae* males to conspecifics than to heterospecific songs from the mainland sister species *F. coelebs* indicates that the song of La Palma chaffinches has diverged to the point where it could act as a reproductive barrier between the two species in case of contact.

However, this inference was not supported for differences between pine and cloud forest birds within *F. canariensis*, suggesting that populations adapted to the two contrasting habitat types may have not yet reached a sufficient degree of lineage divergence or that other pre- or post-mating barriers are at play. As Freeman & Montgomery (2017) highlighted, interpreting playback responses can be challenging: while a lack of response to a song is often taken as evidence of a behavioural barrier to reproduction, a response to a song does not necessarily indicate the absence of such a barrier.

Additionally, playback results do not guarantee that song contributes to reproductive isolation because they primarily test the behaviour of the receiver. For species with closed-ended learning (i.e., species that are not able to modify their repertoire after the first year), like chaffinches, the potential for immigrants to adapt their songs to local dialects is more limited compared to open-ended learners, which have yearly plastic periods (Catchpole & Slater, 2008). This restriction may reduce the likelihood of convergence in song traits across populations and could contribute to maintaining population-specific differences in song. Consequently, this could reinforce the role of song in reproductive isolation, as migrants may be disadvantaged in both territorial disputes and mate attraction if their songs do not align with local dialects. Nevertheless, the effectiveness of song as a barrier will depend on the extent to which population-specific songs are used in mate recognition and whether morphological and ecological differences also contribute to reproductive isolation.

The morphological and genetic adaptations of *F. c. palmae* to cloud and pine habitats suggest that selection has shaped phenotypes to exploit distinct ecological niches (Recuerda *et al*., 2023). For assortative mating to be established or reinforced, a strong linkage between mate recognition traits, such as song, and ecological adaptations is likely critical (Kirkpatrick & Ravigné, 2002). Morphological traits associated with resource exploitation in these two forest types could play a complementary role, as hybrids may have reduced fitness if their intermediate phenotypes are less adapted to either niche. This could impose a cost on immigrants and drive the evolution of assortative mating through reinforcement (Liou & Price, 1994; Coyne & Allen Orr, 1998; Uy *et al*., 2018). At the same time, character displacement in resource-use traits may occur to reduce competition between the two habitat-specific phenotypes, further enhancing ecological divergence (Brown & Wilson, 1956; Schluter & McPhail, 1992).

While our study focused on responses to playback within each forest type, it remains unclear how *F. c. palmae* individuals react in the contact zones between the two forest types. These areas could provide key insights into behavioural isolation, as birds in these zones may exhibit different responses to song. Exploring these ecotones would allow us to determine whether behavioural barriers to reproduction are reinforced, weakened, or altered in transitional environments. Such studies would also help clarify how morphological and ecological divergence interacts with song-based reproductive barriers to influence the evolutionary trajectory of *F. c. palmae* across its range.

## AUTHOR CONTRIBUTIONS

BF: Funding acquisition, Conceptualization, Data curation, Formal analysis, Investigation, Methodology, Project administration, Visualization, Writing – original draft; DG: Conceptualization, Methodology, Investigation, Supervision, Writing – review and editing; CT: Conceptualization, Supervision, Writing – review and editing; BM: Funding acquisition, Conceptualization, Supervision, Project administration, Writing – review and editing

## DATA AVAILABILITY

All the data and code underlying this article are now available as supplementary materials and will be deposited in a public data repository once the article is accepted for publication.

## ACKNOWLEDGEMENTS

We thank Andrea Lirola for her participation in the field recording sessions and for assisting in setting up the initial playback experiments. We also thank all the other field participants that helped recording chaffinches. We thank Max Ringler for his insightful comments, which improved the manuscript. F. Medina, M. Morales and the Cabildo of La Palma provided valuable logistical support during our work on the island. The regional government of the Canary Islands issued the necessary research permits. B.F. was funded by the Foundation for Science and Technology (FCT, Portugal) through a PhD grant (2020.04569.BD; https://doi.org/10.54499/2020.04569.BD). This research was funded by grant PID2021-124501NB-I00 to B.M. from the Spanish Ministry of Science and Innovation, co-financed by the European Union’s Regional Development Fund (ERDF).

